# A Genome-Based Pipeline for Digital Siderophore Typing in *Pseudomonas* Iron-Interaction Network

**DOI:** 10.1101/2025.10.07.680951

**Authors:** Zeyang Qu, Yuanzhe Shao, Anxin Zhang, Lanxin Wang, Ruolin He, Luyan Z. Ma, Shaohua Gu, Zhiyuan Li

## Abstract

Siderophores are key mediators of microbial interactions, especially in iron-limited environments. While traditional experimental methods for siderophore typing are time-consuming and low-throughput, we present a genome-based pipeline for high-throughput digital siderophore typing in *Pseudomonas*. Our approach integrates profile Hidden Markov Models (pHMMs), substrate-specific motif identification, and co-evolutionary analysis to predict both the structure of pyoverdines and their corresponding uptake receptors from any given *Pseudomonas* genome. By this approach, we developed pHMM from 94 previously defined receptor groups and validated their accuracy and robustness across 14,230 *Pseudomonas* genomes. Application of our method to this large dataset demonstrated the potential of our algorithm for identifying both novel “lock-key” receptor groups and previously uncharacterized pyoverdine structures. Notably, our pipeline corrected long-term misclassifications in classical strains and proposed a new reference for the canonical Group III pyoverdine. Furthermore, interaction network analysis supports the observation of distinct siderophore utilization patterns between pathogenic and non-pathogenic strains. This standardized, user-friendly platform offers a robust tool for annotating siderophore behaviors in *Pseudomonas* and demonstrates the potential of digital siderophore typing in exploring iron-mediated ecology across microbes.

**Impact statement:** Iron is vital for microbes. In most environments, they secrete diverse siderophores to scavenge iron. The types of siderophores a strain produces and uptakes shape its interactions, from cooperation to cheating and competition; characterizing this is called “siderophore typing.” While traditional methods are limited by experimental capacities, our genome-based digital pipeline revolutionizes the process by predicting siderophore structures and receptor types directly from genomes, enabling large-scale analysis of 14,230 *Pseudomonas* strains. The results reveal novel groups, correct historical misclassifications, and link patterns to pathogenicity, providing a standardized software for ecological studies, antibiotic design, and bioremediation.

## Introduction

In the microbial world, siderophores mediate social behaviors through iron acquisition[1]. Nearly all microorganisms, with rare exceptions such as *Lactobacillus plantarum*, rely on iron for essential processes, including respiration and DNA biosynthesis[2, 3]. In environments where bioavailable iron is scarce, from oceans (as low as 10^−11^ M)[4] to human tissues (down to 10^−24^ M)[5], siderophores, a diverse class of secondary metabolites, enable microbes to acquire essential iron by chelating Fe^3+^ with remarkable affinity[6]. This iron acquisition process consists of two key steps: synthesis and uptake. Microbes first synthesize and secrete siderophores into the environment, then recognize and transport the resulting siderophore-iron complexes through specific membrane receptors [7, 8]. Beyond simply facilitating individual survival, the ability to produce or utilize specific siderophores influences competitive advantages or cooperative relationships[9]. These interactions are particularly nuanced because siderophores exhibit remarkable structural diversity, with strain-level differences in both production capabilities and uptake specificities[10, 11]. This strain-specific variation creates a complex network where some microbes can exclusively utilize their own siderophores, while others can “pirate” siderophores from other strains[1, 12].

Given this diversity and its profound impact on microbial community dynamics, accurate siderophore typing - the classification of siderophore production and uptake preferences for a strain - has become essential for understanding and predicting microbial behaviors in both natural ecosystems and clinical settings[1, 13]. Despite their critical ecological importance, developing efficient and standardized approaches to perform siderophore typing has remained challenging. Traditional experimental methods for siderophore typing, such as isoelectric focusing[14], mass spectrometry[15], and growth assays[16], have shed valuable light on these molecules but come with notable drawbacks. These techniques often demand painstaking purification of siderophores or receptor proteins, rendering them time-consuming and impractical for analyzing large numbers of samples. What’s more, they frequently blur the line between a strain’s ability to produce a specific siderophore and its capacity to take up various types, a distinction crucial for decoding the game of cooperation and piracy. The lack of consistency across studies and bacterial strains further complicates matters, making it difficult to compare findings or construct a unified framework of siderophore diversity across different species. With genomic data now increasing at an unprecedented rate, there is a pressing need for a faster, more standardized approach, one that can tackle large genomic datasets and clearly distinguish between production and uptake capabilities.

The emergence of genomic-based approaches offers a promising alternative to these experimental constraints. The genus *Pseudomonas* serves as an ideal model for developing such approaches, given its ecological ubiquity, relevance in plant and human diseases, and biotechnological applications[17, 18]. Among *Pseudomonas* siderophores, pyoverdine stands out as particularly well-studied. It is a family of fluorescent green-yellow siderophores synthesized via non-ribosomal peptide synthetase (NRPS) pathways and recognized by specific TonB-dependent membrane receptors called FpvA[18-22]. The landmark work by Jean-Marie et al. in 1997 identified three distinct pyoverdine groups (I, II, and III) in *Pseudomonas aeruginosa* strains and demonstrated these strains could not cross-utilize each other’s pyoverdines, leading to the concept of siderophore typing[13]. By 2024, researchers had experimentally cataloged 79 unique pyoverdine structures[10].

Building on this legacy, our team developed a sequence-to-molecule pipeline that predicts pyoverdine structures directly from genomic data, uncovering 188 unique structures and 94 groups of FpvA receptors across 1,928 *Pseudomonas* genomes, with experimental validation backing its accuracy[23]. Furthermore, our coevolutionary analysis of synthetases and receptors revealed 47 distinct “lock-key” groups. In this system, siderophores produced by the synthetases in each group act as unique keys, specifically recognized and taken up only by the receptors (locks) within their own group, but not by receptors in other groups[24]. This approach allowed us to type siderophores in these 1,928 strains by tracing gene coevolution patterns. However, this “sequence-to-ecology” approach depends on comparing a large set of genomes to detect coevolution, rendering it impractical for typing synthetases and receptors in a single genome, a critical need for detailed studies of individual strains. This gap highlights the urgent demand for a new method tailored to single-genome analysis.

In this study, we introduce the “digital-siderophore-typing” pipeline, which uses our previous genome-coevolution results[23] as a pre-trained model to infer the pyoverdine types produced and taken up by each *Pseudomonas* strain based on its genome. We refined the classification of 94 receptor groups and developed profile Hidden Markov Models (pHMM) for their rapid identification[23]. This computational approach generates a digital representation of siderophore behavior directly from genome annotation files. We validated the pipeline across 14,230 *Pseudomonas* strains, demonstrating its robustness even with fragmented or incomplete genomic data. Moreover, it enables the identification of both novel “lock-key” receptor groups and the determination of previously uncharacterized pyoverdine structures. Testing against Jean-Marie et al.’s landmark strains confirmed accuracy for Groups I and II, while discrepancies in Group III led us to identify an alternative reference strain better suited for future typing efforts. Our analysis revealed distinct siderophore interaction patterns between pathogenic and non-pathogenic strains. To enhance accessibility for experimental biologists, we developed a user-friendly interface that simplifies the typing process. This high-throughput, standardized method establishes a new framework for *Pseudomonas* siderophore typing that can rapidly characterize newly isolated strains and promote consistency across microbiological studies.

## Results

### Workflow for siderophore digital typing and standardized output

Building on our previous work[23], we developed a “sequence-to-function” pipeline for siderophore typing in *Pseudomonas* strains using their genomes. This pipeline identifies the “lock-key groups” for both the pyoverdine synthetase (in producer strains) and all FpvA receptors. It also predicts the molecular structure of the produced pyoverdine based on non-ribosomal peptide synthetase (NRPS) substrate specificity. Together, these analyses enable us to characterize a strain’s “siderophore behavior”—specifically, the type of pyoverdines it can produce, and the types it can uptake (Figure 1A).

**Figure 1.**
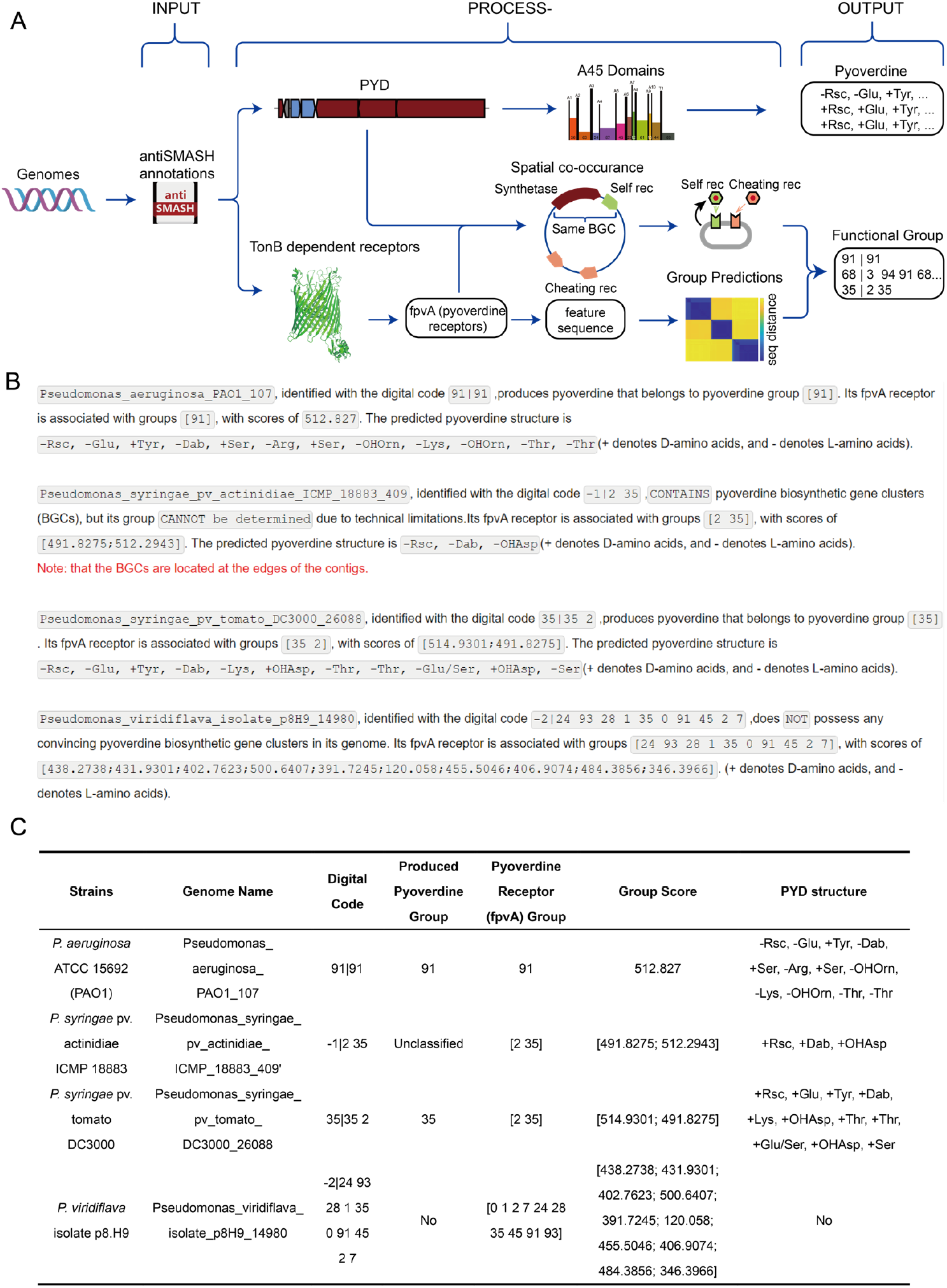
Schematic diagram of the overall workflow for siderophore typing and example outputs. **(A)** Workflow for *Pseudomonas* digital siderophore-typing. The process begins with a genome annotated for secondary metabolites using AntiSMASH, then divides into two parts: identification and annotation of (1) biosynthetic genes and (2) receptor genes. For the first part, pyoverdine biosynthetic genes are initially extracted and located, and their NRPS motifs are extracted to obtain the A45 domain, a key substrate recognition signature sequence. The extracted sequences are aligned with reference A45 domains of known substrates to predict their specificity, which are then merged to predict the complete pyoverdine structure. For the second part, 3 pHMMs (TonB, STN, and Plug domains) are used to extract all potential TonB-dependent receptors, then two characteristic sequences are used to identify the pyoverdine recognition receptor (fpvA). The fpvA sequences undergo feature sequence extraction[23] and are matched against 94 existing pHMMs to classify their corresponding pyoverdine groups. Subsequently, the co-occurrence of pyoverdine biosynthetic genes and receptor genes within the same biosynthetic gene cluster (BGC) distinguishes self-receptors (for uptake of self-produced pyoverdine) from cheating receptors (for uptake of heterologous pyoverdine). This workflow elucidates the siderophore behavior of *Pseudomonas*, including the types of pyoverdine it can produce and uptake, and the predicted structure of the produced pyoverdine. **(B)** Example textual output of the siderophore typing procedure. **(C)** Tabular summary of siderophore typing results. The output encompasses the input genome name, a digitized representation of its pyoverdine siderophore behavior, the classification of the produced pyoverdine group (determined via the self-receptor), the classification of the produced pyoverdine receptor (fpvA) group, along with prediction scores for these receptor groups, and the predicted pyoverdine structure.

The workflow begins with genome annotation using antiSMASH (version 7.0.0 or higher recommended)[25] to identify secondary metabolite gene clusters. It then splits into two parallel paths:

#### Path 1: Molecular Structure Prediction (Figure 1A, upper path)

- Pyoverdine-synthesizing gene sequences are extracted from the antiSMASH-annotated genome file.
- These sequences are assembled into a standard NRPS structure.
- The A45 domain—a crucial substrate recognition sequence from the non-ribosomal peptide synthetase (NRPS)[26] was identified by the NRPSMotifFinder[26].
- The previously proposed “phylogeny-focused approach” is applied to the A45 domain to predict the complete pyoverdine[23].

#### Path 2: Annotation of the “lock-key” groups for all fpvA receptors

- All TonB-dependent receptors (TonBDR) in the genomes are annotated by profile Hidden Markov Models (pHMM) selecting for the three typical TonBDR domains (TonB, STN, and Plug).
- The pyoverdine recognition receptors, FpvA, among all TonBDRs, are pinpointed by two key sequence regions from our earlier work[23].
- A highly discriminatory “feature sequence” (near the Plug domain, from 168 Pro to 295 Ala in PAO1) is extracted from each fpvA sequence, which has been demonstrated to effectively distinguish different “lock-key” groups[23].
- Each feature sequence is matched against 94 predefined pHMMs. These pHMMs are constructed using feature sequences from 94 distinct groups of 4,547 FpvA receptors, which have been previously identified across 1,928 *Pseudomonas* genomes[23]. This matching process predicts the groups of pyoverdine that the strain can uptake.
- “Self-receptor” is defined as the receptor that uptakes the siderophore produced by the strain itself. Based on prior observations that self-receptors are typically co-located with pyoverdine biosynthetic genes within the same biosynthetic gene cluster (BGC) [24, 27-29], we designate receptors encoded within the pyoverdine BGC as self-receptors. Receptors located outside this BGC are classified as “cheating” receptors, which may recognize pyoverdines produced by other strains.

### Standardized Output

After completing the above steps, the pipeline generates a standardized textual output consisting of six elements for each strain:

- **Genome name**: The identifier of the input strain.
- **Digital code for siderophore function groups**: A numerical representation of the strain’s siderophore behavior, formatted as “self-receptor group | all receptor groups.” The part before “|” indicates the pyoverdine type produced (via the self-receptor), while the part after “|” lists all receptor groups, reflecting the pyoverdine types the strain can uptake.
- **Produced pyoverdine group**: Determined by the self-receptor group.
- **FpvA receptor groups**: The classified groups of all FpvA receptors in the strain.
- **Score for each FpvA receptor group**: HMM scores reflecting the similarity between the strain’s receptors and their assigned groups.
- **Pyoverdine molecular structure**: A predicted sequence of concatenated monomer units representing the synthesized pyoverdine, where “+” denotes the D-configuration and “−” denotes the L-configuration.

To demonstrate the pipeline’s application, we present the output for four *Pseudomonas* strains with distinct siderophore behaviors: *P. aeruginosa* ATCC 15692 (PAO1), *P. aeruginosa* ATCC 27853, *P. syringae* pv. tomato DC3000, and *P. viridiflava* isolate p8.H9 (Figure 1B), and summarize the results in the table shown in Figure 1C.

### Distribution of Pyoverdine Groups in 14,230 *Pseudomonas* Genomes

We tested the current pipeline on all 14,230 *Pseudomonas* genomes available in existing *Pseudomonas* databases[30], validating the robustness of our processing workflow. Next, we analyzed the distribution of different pyoverdine groups across these genomes. For the classification of synthesized pyoverdines, we use the self-receptor group as a proxy for the produced pyoverdine group. As shown in Figure 2A, the most prevalent groups included the three classical pyoverdine types—Group I, II, and III—originally identified by Jean-Marie Meyer et al. in *Pseudomonas aeruginosa*[13], which is likely due to the high representation of *P. aeruginosa* in current sequencing datasets. Additionally, we observed a high proportion of Group 35 pyoverdine, a siderophore type widely present in plant-associated *Pseudomonas* species, such as *Pseudomonas syringae*. For the ten most abundant pyoverdine groups, we provide representative structures, identified as the most frequently predicted structures within each group, along with representative strains.

**Figure 2.**
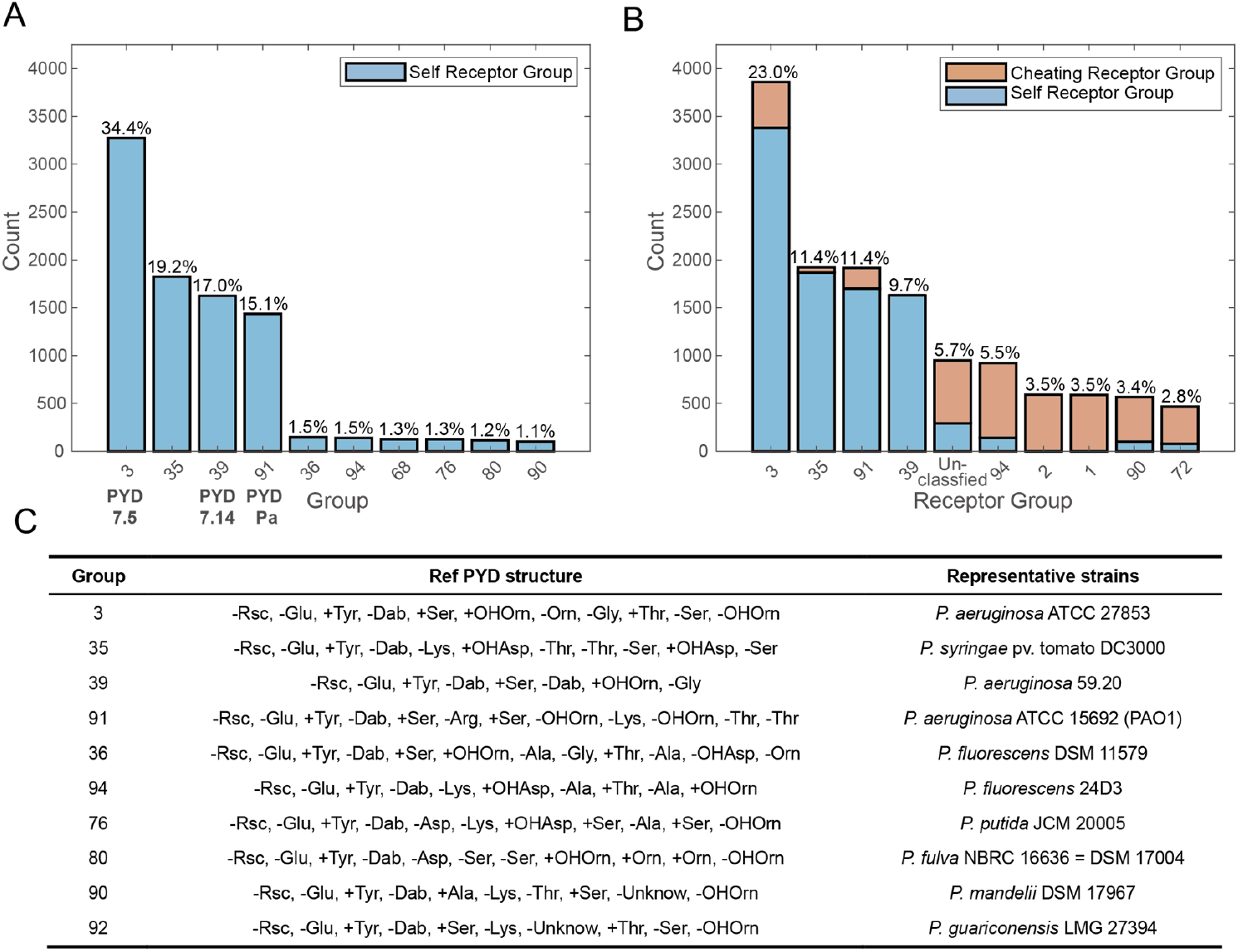
Distribution of Siderophore Pyoverdine Behavior in the *Pseudomonas* Genus from the *Pseudomonas* Database. **(A)** Distribution of pyoverdine (PVD) biosynthesis groups within the *Pseudomonas* genus, classified based on self-receptor recognition. Only the 10 most abundant groups are shown. **(B)** Distribution of pyoverdine receptor (fpvA) groups in the *Pseudomonas* genus, displaying the 10 most prevalent groups. Bars are color-coded: the blue portion represents the proportion of self-receptors, while the orange portion denotes cheating receptors. **(C)** Representative pyoverdine structures and corresponding *Pseudomonas* strains for the 10 most abundant biosynthetic groups. The structures shown here represent the most commonly predicted structures within each group.

In the analysis of pyoverdine uptake capabilities, we identified the ten most prevalent receptor groups (Figure 2B), and unsurprisingly, the distribution is similar to that of synthetase. We also distinguished self-receptors (which uptake the strain’s own pyoverdine) from cheating receptors (which uptake pyoverdines produced by other strains), and noticed that less prevalent receptor groups generally exhibited a higher proportion of cheating receptors. Notably, 5.7% of receptors could not be assigned to any of the 94 predicted groups based on our current algorithm.

As previously defined, we conveniently classify receptors encoded within the pyoverdine biosynthetic gene cluster (BGC) as “self-receptors” and those outside the BGC as “cheating receptors”. To validate this, we analyzed lock-key relationships in 8,701 filtered pyoverdine structure records from 14,230 *Pseudomonas* genomes using two complementary approaches and found high consistency (Figure S1). The first approach matches the BGC-encoded receptor gene against 94 predefined pHMMs to identify the self-receptor group. The second predicts the pyoverdine structure directly from BGC synthetase sequences by their A domains, then compares it to 183 reference structures in 1,928 representative strains[24]; each reference links to a specific self-receptor group via lock-key relationships, enabling group assignment. For exact matches (4,919 records), the two approaches agreed in 99.88% of cases. For non-exact matches (3,782 records), we identified the closest reference using the longest common subsequence and minimal edit distance, yielding around 90% consistency. These results confirm the reliability of BGC-encoded receptors for identifying and grouping self-receptors.

### Discovery of new pyoverdine groups and determining the structure of known groups

In addition to annotating known siderophore synthetase and receptors, this approach also demonstrates strong potential for discovering novel lock-key groups. In our large-scale analysis of 8,701 pyoverdine structures from 14,230 *Pseudomonas* genomes, we identified three potential new lock-key groups (designated as Groups 95, 96, and 97, Figure 3A). These newly discovered groups exhibit substantial divergence from the 94 established reference groups[23] in both pyoverdine structure and receptor sequence, confirming their novelty. Upon comparison with the SIDERITE database[10], we found that one of these structures had been recorded in this database (Group 95) but was not included in our previous work[23], while the remaining two structures appear to be entirely novel (Group 96 and Group 97).

**Figure 3.**
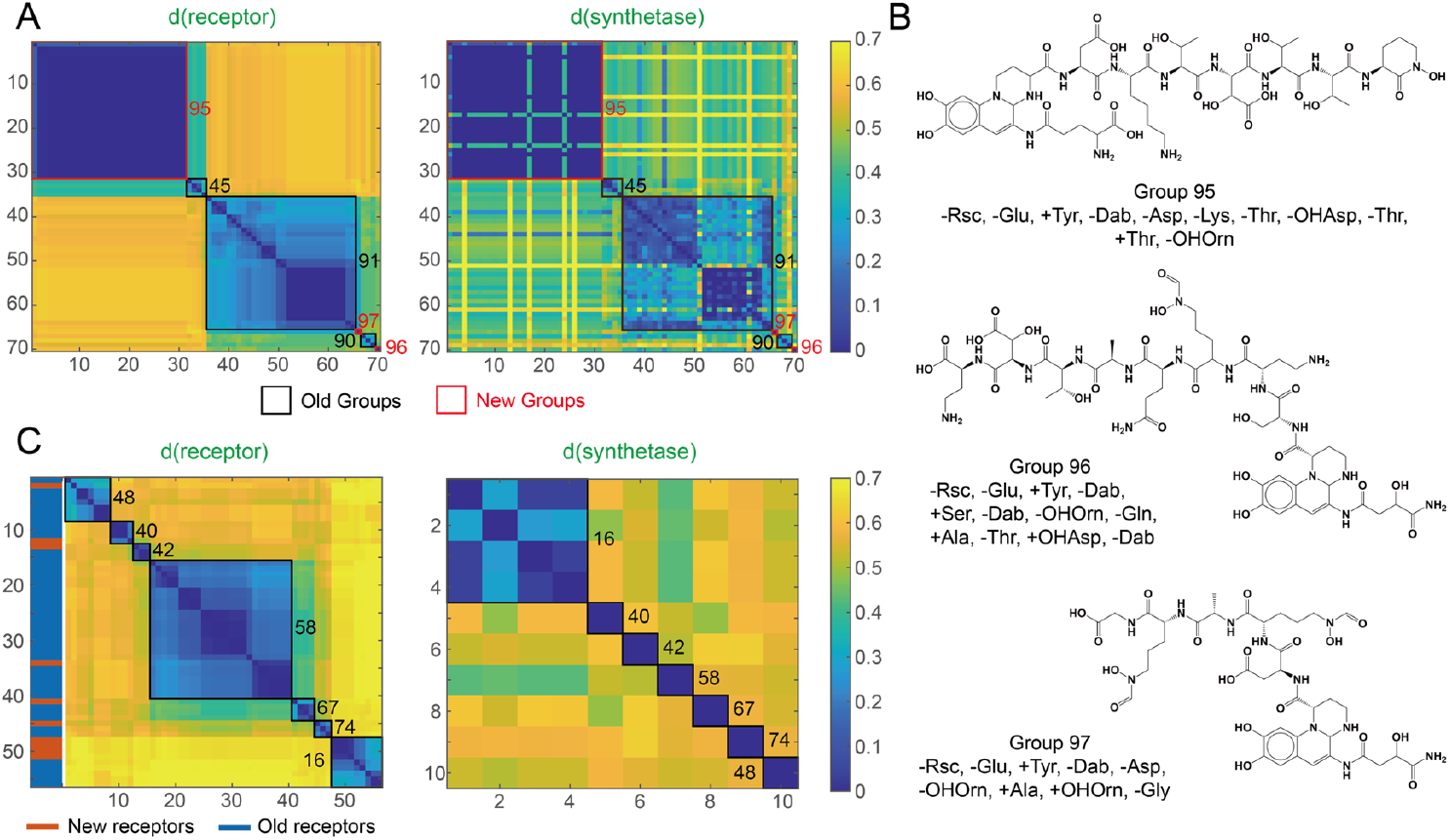
Identification of new pyoverdine groups or confirmation of known groups using the siderophore typing approach. (A) Heatmaps showing the sequence distances of receptors (left) and biosynthetic genes (right) between newly identified pyoverdine groups and their closest matches among the previously defined 94 groups[23]. Red boxes indicate the newly discovered groups (Groups 95, 96, and 97), while black boxes represent the most similar groups among the existing 94(Groups 45, 90, and 91, respectively), based on receptor gene sequences. Receptor gene distances are calculated using the feature sequences of the receptor genes (Left panel)[23]. Biosynthetic gene distances are computed using the alignment approach developed in our previous study (right panel)[23]. The results show that these newly identified groups differ substantially from previously characterized groups at both the receptor and biosynthetic gene levels. (B) The chemical structure of Group 95 was retrieved from the SIDERITE database[10], whereas the structures of Groups 96 and 97 were constructed by linking the predicted amino acid sequences to the chromophore through peptide bonds and were visualized using ChemDraw. (C) Prediction of pyoverdine structures for known groups across a broader set of genomes. The sequence distances of receptor genes for Groups 16, 40, 42, 48, 58, 67, and 74 are shown in the left panel. Blue bars represent receptors identified in our previous study[23], while orange bars indicate novel receptors that were found to be associated with corresponding biosynthetic genes in this study. The right panel shows the sequence distances among the biosynthetic genes of these same groups, calculated using the p-distance method.

Previously, due to limitations in the dataset size, we were only able to predict the receptor group for many pyoverdine types without identifying the corresponding biosynthetic gene sequences[24], leaving some “unmatched groups”. In this study, the expanded dataset allowed us to address this limitation by identifying pyoverdine structures for several previously “unmatched groups”, including Groups 16, 40, 42, 48, 58, 67, and 74, for which no associated synthetase genes or structural information had been found previously[23] (Figure 3C). Sequence alignment revealed that the self-receptors linked to these newly identified structures are highly similar to the known receptors of their respective groups. However, the corresponding pyoverdine structures differ substantially from those previously described, supporting the assignment of these structures to the groups that had lacked defined structures before. These findings further highlight the effectiveness of our method as a convenient and efficient tool for identifying both novel lock-key receptor groups and previously uncharacterized pyoverdine structures.

### Reassessment of the Classical Pyoverdine Group and Reclassification of the Group III Pyoverdines

Subsequently, we applied our algorithm to analyze the genomes of classical strains associated with Group I, II, and III pyoverdines, including *P. aeruginosa* ATCC 15692 (PAO1), *P. aeruginosa* ATCC 27853, *P. aeruginosa* Pa.6, and *P. aeruginosa* 59.20[13]. Our analysis revealed certain discrepancies with previous understanding: For *P. aeruginosa* ATCC 15692 (PAO1), *P. aeruginosa* ATCC 27853, and *P. aeruginosa* 59.20, our findings align with prior reports, confirming their correspondence to Group I, Group II, and Group III pyoverdines, respectively. However, in the case of *P. aeruginosa* Pa.6, which has been documented as a typical strain to produce Group III pyoverdine, our genomic predictions indicate that this strain actually produces Group II pyoverdine. Further examination revealed that *P. aeruginosa* Pa.6 shares a similar pyoverdine structure with *P. aeruginosa* ATCC 27853, and its receptor belongs to the same group as that of *P. aeruginosa* ATCC 27853. Based on these observations, we hypothesize that strain variations in *P. aeruginosa* Pa.6, potentially introduced during laboratory passage and sequencing, may account for this inconsistency.

Therefore, to accurately identify strains capable of producing Group III pyoverdine, we used our pipeline to screen for candidates. We discovered that *Pseudomonas* IMP66[31], a strain isolated from crude oil, was predicted to produce Group III pyoverdine based on both its self-receptor classification and structural analysis. Experiments confirmed our prediction: *Pseudomonads* IMP66 produce a high concentration of pyoverdine (Figure 3 B), and mass spectrometry analysis validated that *Pseudomonads* IMP66 did not produce Group I or II pyoverdines but exclusively synthesized pyoverdine of the same signature of Group III (Figure 3 C). Thus, this strain has the potential to replace *P. aeruginosa* Pa.6 as the model strain for studying Group III pyoverdine.

**Figure 3.**
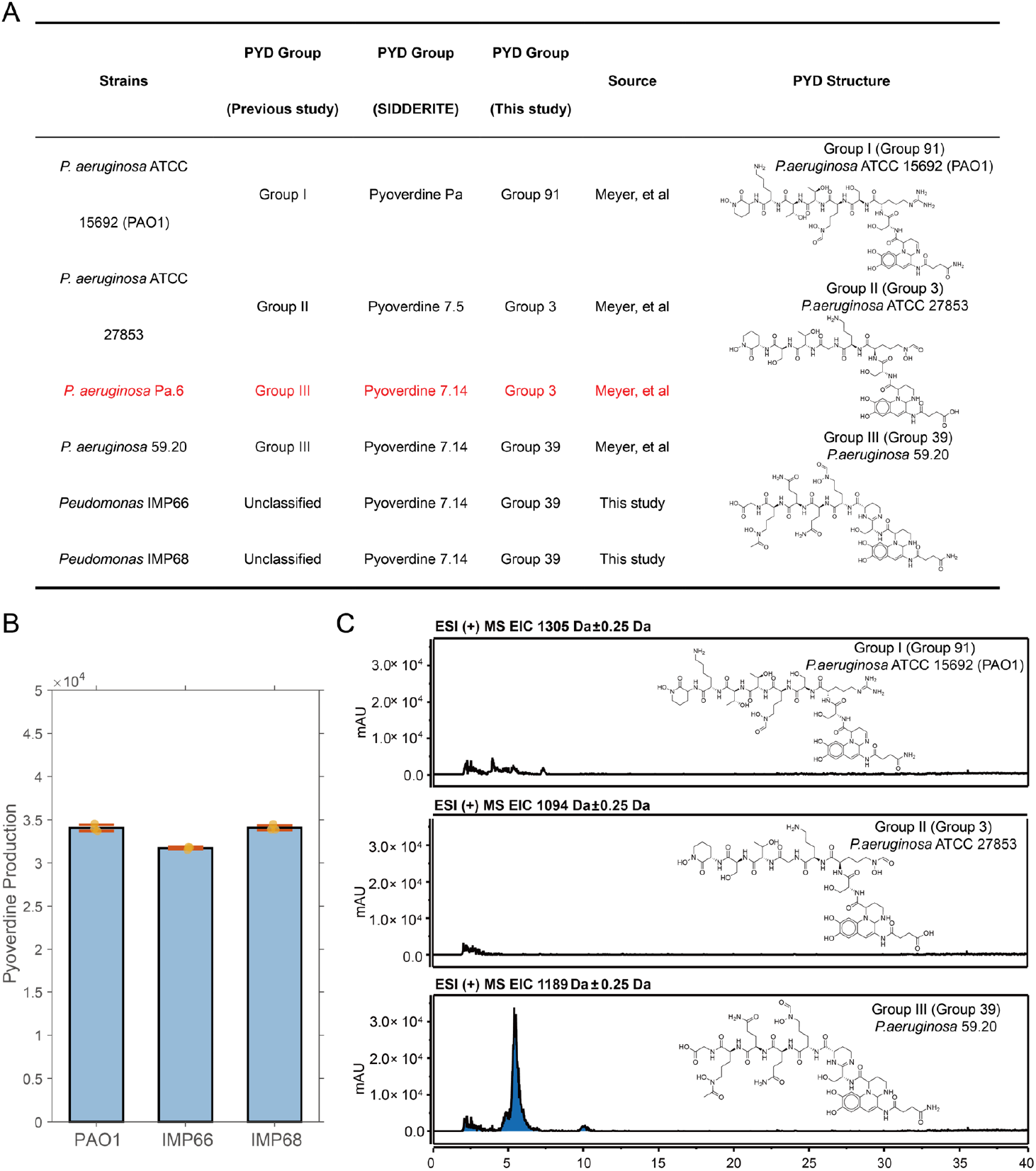
Reevaluation and Reclassification of Group III Siderophores. **(A)** Three classical siderophore groups and their corresponding *Pseudomona*s strains, as identified in the work of Meyer et al. In this study, the siderophore group classification of these strains, along with five additional strains, was re-evaluated. Discrepancies between the original classification and the reclassification in this study are highlighted in red. **(B)** Comparison of siderophore production between *Pseudomonas* IMP66 and *P. aeruginosa* PAO1, measured by fluorescence intensity at wavelengths of 398–460 nm. **(C)** Mass spectrometry (MS) analysis of siderophores produced by *Pseudomonas* IMP66. The extracted ion chromatograms (EIC) display the molecular masses corresponding to pyoverdines of Group I, Group II, and Group III from top to bottom, respectively.

### Digital Siderophore Typing Strengthens the Link Between Pathogenicity and Pyoverdine Utilization Strategies

After analyzing all 14,230 *Pseudomonas* strains from the *Pseudomonas* Database (Figure 2), we constructed the siderophore interaction network encompassing all strains (Figure 4A), more comprehensive than the previous 1982-strain network [24]. We then extracted and examined the siderophore interaction networks within several pathogenic and non-pathogenic *Pseudomonas* species separately (Figure 4 B). Our findings agreed that the siderophore interaction network in pathogenic species, such as *P. aeruginosa* and *P. syringae*, is markedly simpler compared to that of non-pathogenic species, such as *P. fluorescens* and *P. putida*. Moreover, pathogenic strains predominantly exhibit pyoverdine behaviors characterized by single-receptor producers and nonproducers, whereas non-pathogenic strains primarily function as multi-receptor producers. This pattern aligns with the previous findings from a smaller dataset of 1,928 representative *Pseudomonas* strains, further reinforcing the connection between pathogenicity and pyoverdine utilization strategies[24].

**Figure 4.**
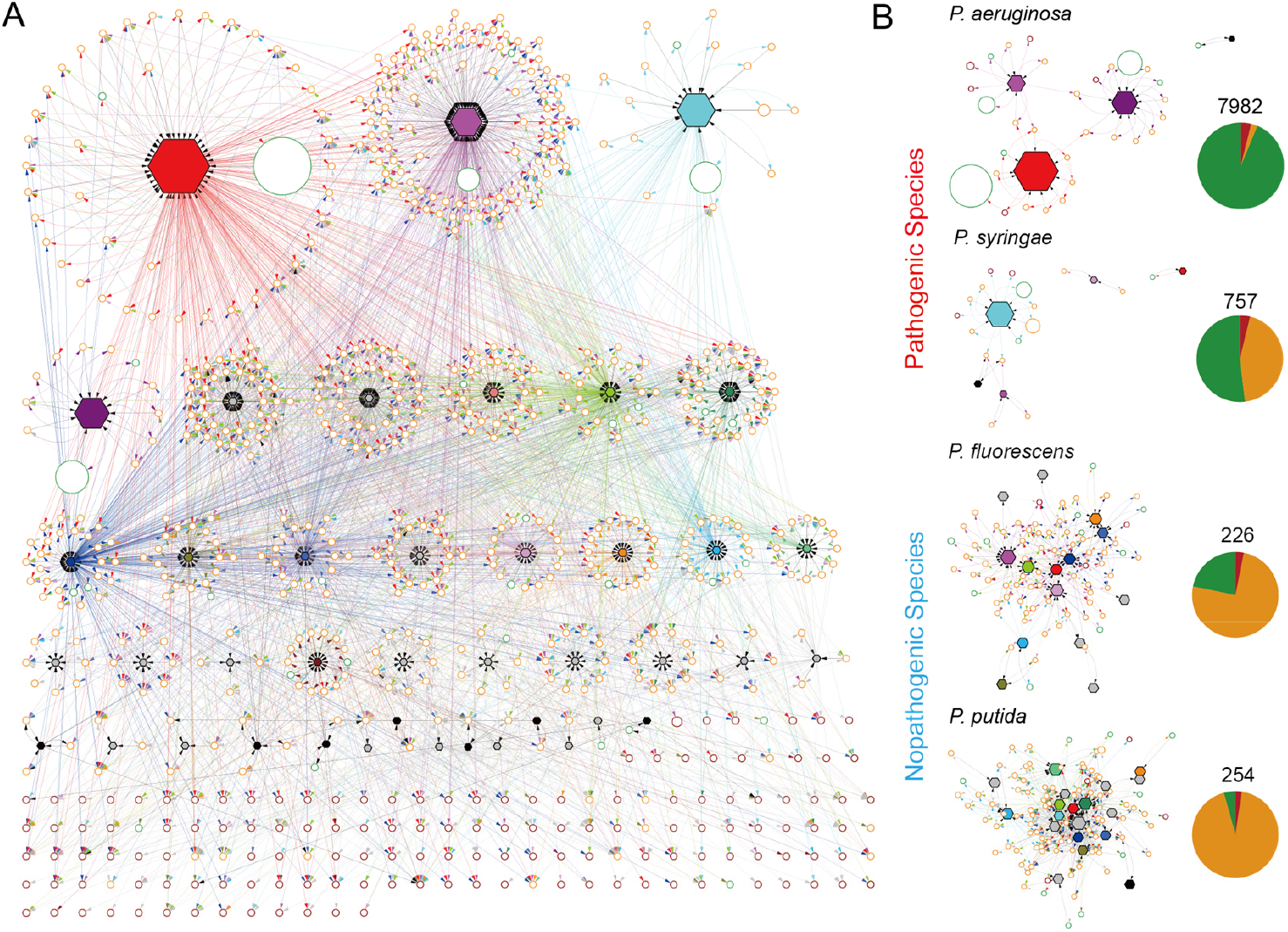
Pyoverdine-mediated interaction network among 14,230 *Pseudomonas* strains from the *Pseudomonas* Database. **(A)** The predicted siderophore interaction networks facilitated by pyoverdines among all 14,230 *Pseudomonas* strains. Circular nodes represent functional groups, defined as strains that produce the same pyoverdine type and utilize an identical set of pyoverdines. The size of each circular node is proportional to the number of strains within the corresponding functional group. Circular node colors indicate single-receptor producers (green), multi-receptor producers (yellow), and nonproducers (red)[24]. Hexagonal nodes represent distinct pyoverdine groups, with node size corresponding to the number of strains utilizing that particular pyoverdine. The pyoverdine groups exclusively associated with single-receptor producers are highlighted in their respective receptor group colors, whereas pyoverdine groups found exclusively in multi-receptor producers are shown in gray. Directed edges from circular to hexagonal nodes indicate pyoverdine production, while edges from hexagonal to circular nodes denote pyoverdine utilization, with edge colors matching the functional group classification. **(B)** Pyoverdine-mediated iron interaction network across pathogenic and nonpathogenic *Pseudomonas* species.

## Discussion

Siderophore typing, the classification of *Pseudomonas* strains based on the types of pyoverdines they produce and uptake, has become a classical method to characterize *Pseudomonas* iron scavenging behavior. In this study, we addressed three key challenges. First, traditional *Pseudomonas* siderophore typing methods—including isoelectric focusing (IEF), mass spectrometry (MS), ion mobility spectrometry, ferripyoverdine outer-membrane receptor immunoblotting, iron uptake assays, and growth stimulation assays—rely on experimental techniques that are inherently low-throughput, unable to simultaneously determine both pyoverdine production and uptake profiles, and difficult to compare across different methods[32, 33]. Our genome-based approach overcomes these limitations by offering a standardized, high-throughput, and integrated siderophore typing method, with a user-friendly graphical interface that facilitates widespread adoption in the field. Second, we provided the first large-scale analysis of siderophore behavior across all publicly sequenced *Pseudomonas* strains, offering a valuable resource for *Pseudomonas* research. Third, we identified potential inaccuracies in the reference strain for Group III pyoverdine production and established a new standard strain, suggesting that previous studies using *P. aeruginosa* Pa.6 should be reevaluated.

The discovery of novel receptor groups and the characterization of synthetases for previously unmatched groups[23] in our expanded dataset further underscores the value of large-scale genomic analyses, highlighting the need for continued microbial sequencing efforts to uncover hidden siderophore diversity. For pathogenic species like *P. aeruginosa*, this precise understanding enables targeted interventions, such as siderophore-antibiotic conjugates that exploit iron acquisition pathways to enhance antimicrobial efficacy[34-36]. In contrast, the siderophore diversity in non-pathogenic strains holds promising applications in agriculture, including biofertilizers that improve plant iron uptake in deficient soils and in bioremediation of heavy metal-contaminated environments through chelation[37]. However, given the strain-level difference of the iron uptake pathways, effective interventions demand accurate characterization of the siderophore types produced and utilized by the target strains to avoid inadvertently aiding pathogens or disrupting beneficial probiotics’ iron uptake.

Another interesting insight is that classical microbiological classifications are vulnerable to decades of accumulated errors, as demonstrated by the *P. aeruginosa* Pa.6 misclassification likely caused by laboratory passage or sequencing artifacts. Genomic reassessments have revealed similar inconsistencies in reference strains PAO1 and PA14, where core genome analyses identify environmental adaptations missed by traditional phenotyping[38, 39]. Such misclassifications may have compromised studies of virulence, antibiotic resistance, and ecological dynamics, making genomic reevaluation essential. Our digital typing may help to address this challenge by enabling rapid, scalable strain verification directly from sequence data, again demonstrating that larger datasets yield more comprehensive insights.

Despite its strengths, our pipeline has limitations: it relies on high-quality genome assemblies, potentially underperforming on fragmented data, and it focuses solely on pyoverdines, excluding other siderophores like pyochelin[40]. Database biases toward *P. aeruginosa* may overlook underrepresented species. Also, while predictive accuracy is high (e.g., 99.88% consistency), experimental validation remains essential for novel predictions. Future extensions could incorporate machine learning for broader siderophore classes and genera, integrating tools with emerging databases for comprehensive metallophore mining[41]. In conclusion, this work marks a paradigm shift from labor-intensive experimental typing to digital, genomics-driven approaches, fostering consistency and discovery in siderophore ecology. By resolving historical inaccuracies and unveiling new diversity, it paves the way for predictive microbiology in the big data era, with applications spanning infection control to sustainable agriculture. Expanding beyond *Pseudomonas*, similar frameworks could revolutionize iron acquisition studies across microbes, ultimately advancing our understanding of microbial evolution and interactions.

## Material and methods

The installer for this platform is readily accessible via https://sidtyping2025-1342626677.cos.ap-beijing.myqcloud.com/Installation/SiderophoreTyping_Setup.zip, facilitating a straightforward installation process.

## Acknowledgment

This work was supported by the National Key Research and Development Program of China (No. 2024YFA0919500) and the National Natural Science Foundation of China (No. 42577142 and No. T2321001). LZ was supported in part by the Peking-Tsinghua Center for Life Sciences.

## Supplementary Figure

**Figure S1.**
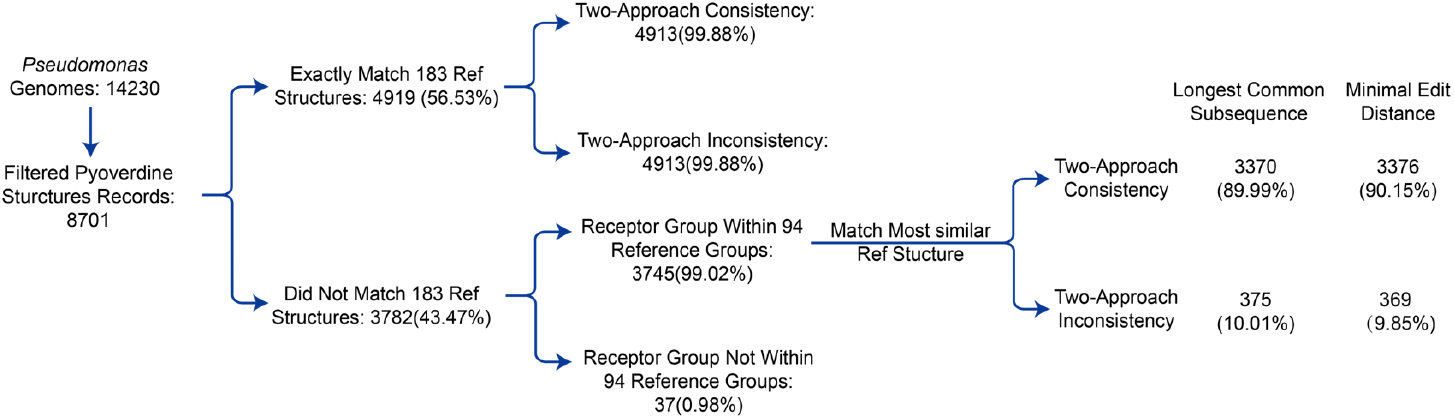
Figure S1. Validation of self-receptor classification in *Pseudomonas* pyoverdine biosynthetic gene clusters (BGCs). Flowchart summarizing the consistency analysis of two complementary methods for predicting self-receptors from 8,701 pyoverdine structure records derived from 14,230 *Pseudomonas* genomes. The first approach matches BGC-encoded receptor genes to 94 predefined pHMMs, while the second infers pyoverdine structures from BGC synthetase genes and compares them to 183 reference structures. Among records exactly matching references (4,919/8,701, 56.53%), self-receptor assignments showed 99.88% consistency between methods. For non-matching structures (3,782 records; 43.47%), the overall approach remained unchanged, while the second method, which uses the closest reference structure to the query based on the longest common subsequence or minimal edit distance, achieved consistent receptor group assignments in 89.99% and 90.15% of cases, respectively.

